# Simultaneous suppression of ribosome biogenesis and Tor activation by TRIM-NHL proteins promotes terminal differentiation

**DOI:** 10.1101/2021.03.31.437822

**Authors:** Jinghua Gui, Tamsin J Samuels, Katarina ZA Grobicki, Felipe Karam Teixeira

## Abstract

Proper tissue development and homeostasis depends on the balance between growth and terminal differentiation, but the mechanisms coordinating these processes remain elusive. Accumulating evidence indicates that ribosome biogenesis (RiBi) and protein synthesis, two of the most energy-consuming cellular processes sustaining growth, are tightly regulated and yet can be uncoupled during stem cell fate transitions. Here, using the *Drosophila* adult female germline stem cell (GSC) and larval neuroblast (NB) systems, we show that Mei-P26 and Brat, two *Drosophila* TRIM-NHL paralogs of the mammalian TRIM32 protein family, are responsible for uncoupling RiBi and protein synthesis during GSC and NB differentiation, respectively. Mei-P26 and Brat modify the metabolism of differentiating cells by activating the Target of rapamycin (Tor) kinase to promote translation, while concomitantly repressing RiBi. Depletion of Mei-P26 or Brat results in excessive cellular growth and defective terminal differentiation, which can be rescued by ectopic activation of Tor together with suppression of RiBi. Our results indicate that the anabolic reprogramming established by TRIM-NHL activity by uncoupling RiBi and translation activities creates the conditions required for terminal differentiation.

## Introduction

The tight coordination of cellular metabolic activities is essential for cell growth, proliferation, stress response, and cell survival, and therefore is at the core of the developmental processes sculpting complex organs and enabling robust tissue repair and homeostasis in adults (Buszczak et al., 2014; Chua et al., 2020; Miyazawa & Aulehla, 2018). For instance, translation and ribosome biogenesis (RiBi), two of the most energy-consuming anabolic activities, have been shown to be dynamically regulated during cell fate transitions, with stem cells differing significantly from their immediate differentiating progeny (Teixeira & Lehmann, 2019). This dynamic regulation has been observed in many stem cell systems, including the *Drosophila* adult germline stem cells, intestinal stem cells, and larval neuroblasts (Betschinger et al., 2006; Hung et al., 2020; Neumüller et al., 2008; Sanchez et al., 2016; Zhang et al., 2014), as well as a variety of mouse adult stem cells in the hematopoietic, neural, muscle, and skin systems (Baser et al., 2019; Blanco et al., 2016; Llorens-Bobadilla et al., 2015; Sampath et al., 2008; Signer et al., 2014; Zismanov et al., 2016). Genetic and pharmacological manipulations revealed that the control of these metabolic activities is critically important for tissue homeostasis, tipping the balance between self-renewal and differentiation (Baser et al., 2019; Blanco et al., 2016; Buszczak et al., 2014; Sanchez et al., 2016; Signer et al., 2014). However, even though dynamic changes in translation and RiBi are required during stem cell differentiation and are pervasive across different systems, the mechanisms by which they are regulated remain poorly understood.

The *Drosophila* ovary presents an ideal *in vivo* system for dissecting the regulation of metabolism during stem cell differentiation (Spradling et al., 2011). Germline stem cells (GSCs), found attached to the somatic niche at the anterior-most part of the ovaries, show lower translational activity and higher RiBi rates in comparison to neighboring differentiating cells (Neumüller et al., 2008; Sanchez et al., 2016; Zhang et al., 2014). Upon stem cell division and niche exclusion, the differentiating cystoblast (CB) undergoes four rounds of mitosis with incomplete cytokinesis before terminally differentiating as a 16-cell cyst. The differentiating cyst stages are characterized by higher translation and lower RiBi compared to the stem cell (Supplementary figure 1A, (Sanchez et al., 2016)), which is thought to result in limited production of new ribosomes alongside high demand for protein synthesis. Indeed, translation and RiBi are usually closely coordinated to ensure adequate numbers of ribosomes are available to sustain the required levels of protein synthesis (G. Y. Liu & Sabatini, 2020). Therefore it is not surprising that the metabolic changes observed during GSC differentiation are associated with changes in growth, with differentiating progeny decreasing in cell size prior to terminal differentiation (Neumüller et al., 2008). Notably, experimental modulation of RiBi and protein synthesis activities during germline differentiation has been shown to affect the balance between self-renewal and differentiation, resulting in either premature loss of GSCs, a block in differentiation, or tumorigenesis (Neumüller et al., 2008; Sanchez et al., 2016; Sun et al., 2010; Zhang et al., 2014).

A known regulator of RiBi during GSC differentiation is Meiotic P26 (Mei-P26), a germline-expressed gene encoding a member of the evolutionarily conserved TRIM-NHL family of proteins (the mammalian TRIM32 family) (Neumüller et al., 2008; Page et al., 2000). In *mei-P26* mutant ovaries, differentiating cysts maintain high RiBi rates that are usually characteristic of stem cells, overgrow, and fail to terminally differentiate, leading to the formation of tumors composed of partially differentiated cysts (Neumüller et al., 2008; Page et al., 2000). Similarly, in the *Drosophila* larval brain, mutants of the *mei-P26* paralog *brain tumor* (*brat*) are characterized by the failure of differentiation of the progeny of the larval neuroblasts (NBs). *brat* mutant progeny cells show increased RiBi rates, large cell size and excessive proliferation – leading to a characteristic brain tumor phenotype after which the gene was originally named (Betschinger et al., 2006; Lee et al., 2006).

Here, we delve into the mechanisms that control protein synthesis and RiBi activity during stem cell differentiation. First, we show that the activity of the Target of rapamycin (Tor) kinase – an evolutionarily conserved regulator of cell metabolism that generally coordinates translation and RiBi to drive growth -is developmentally regulated during stem cell differentiation. We demonstrate that Tor activation drives the observed increase in protein synthesis during germline differentiation. Surprisingly, we find that Mei-P26 and Brat are activators of the Tor kinase, alongside their previously identified roles in suppressing RiBi. While *mei-P26* and *brat* mutant cells do not activate the Tor kinase during differentiation, overexpression of these TRIM-NHL proteins leads to ectopic Tor activation, which results in premature differentiation. Using genetic and pharmacological manipulations, we show that the *mei-P26-*or *brat-*induced block in differentiation can be attenuated by restoring RiBi suppression and Tor activation, revealing that the metabolic uncoupling driven by TRIM-NHL proteins is critical for inducing terminal differentiation.

## Results

### Tor is activated during GSC differentiation and is required for the increase in translation rate

We have previously shown that RiBi and protein synthesis rates are actively regulated and yet uncoupled during GSC differentiation, when protein synthesis increases alongside a reduction in RiBi (Sanchez et al., 2016). This uncoupling is at odds with the well-established role of the evolutionary conserved Tor kinase in coordinating RiBi and protein synthesis activities to promote growth (G. Y. Liu & Sabatini, 2020). Tor kinase activity has been previously shown to be involved in GSC proliferation and cyst growth (LaFever et al., 2010; Sanchez et al., 2016; Sun et al., 2010) but the loss of Tor activity does not affect RiBi in GSCs (Sanchez et al., 2016), raising the question of when the Tor kinase is active during GSC differentiation. To investigate this, we took advantage of an antibody against the phosphorylated form of the ribosomal protein S6 (p-S6), a downstream target and readout of the activity of the Tor pathway (Pullen et al., 1998; Romero-Pozuelo et al., 2017).

Immunofluorescence microscopy analysis revealed that GSCs were devoid of p-S6, but strong p-S6 signal was detectable from the differentiating CB stage onwards (Figure 1A, B, Supplementary figure 1A). While only ∼12% of CBs were positively marked by p-S6, all 2-cell and ∼93% of 4-cell cysts were p-S6^+^, with the penetrance of p-S6 signal declining in 8-cell (∼59%) and terminally differentiated 16-cell cysts (∼11%). Analysis using the cell cycle tracing FUCCI system (Zielke et al., 2014) confirmed that p-S6 expression was not detected in GSCs regardless of the cell-cycle phase, but was present in cysts throughout the cell cycle (Supplementary figure 1B-D). Moreover, a short incubation of adult ovaries with rapamycin, a specific inhibitor of the Tor kinase, was sufficient to abolish the p-S6 signal in differentiating cells (Figure 1C). As the Tor kinase and the Target of Rapamycin Complex 1 (TORC1) co-factor Raptor proteins are present from GSCs to 16-cell cysts (Supplementary figure 1E), our results indicate that Tor is inactive in the GSC but is activated during differentiation.

**Figure 1.**
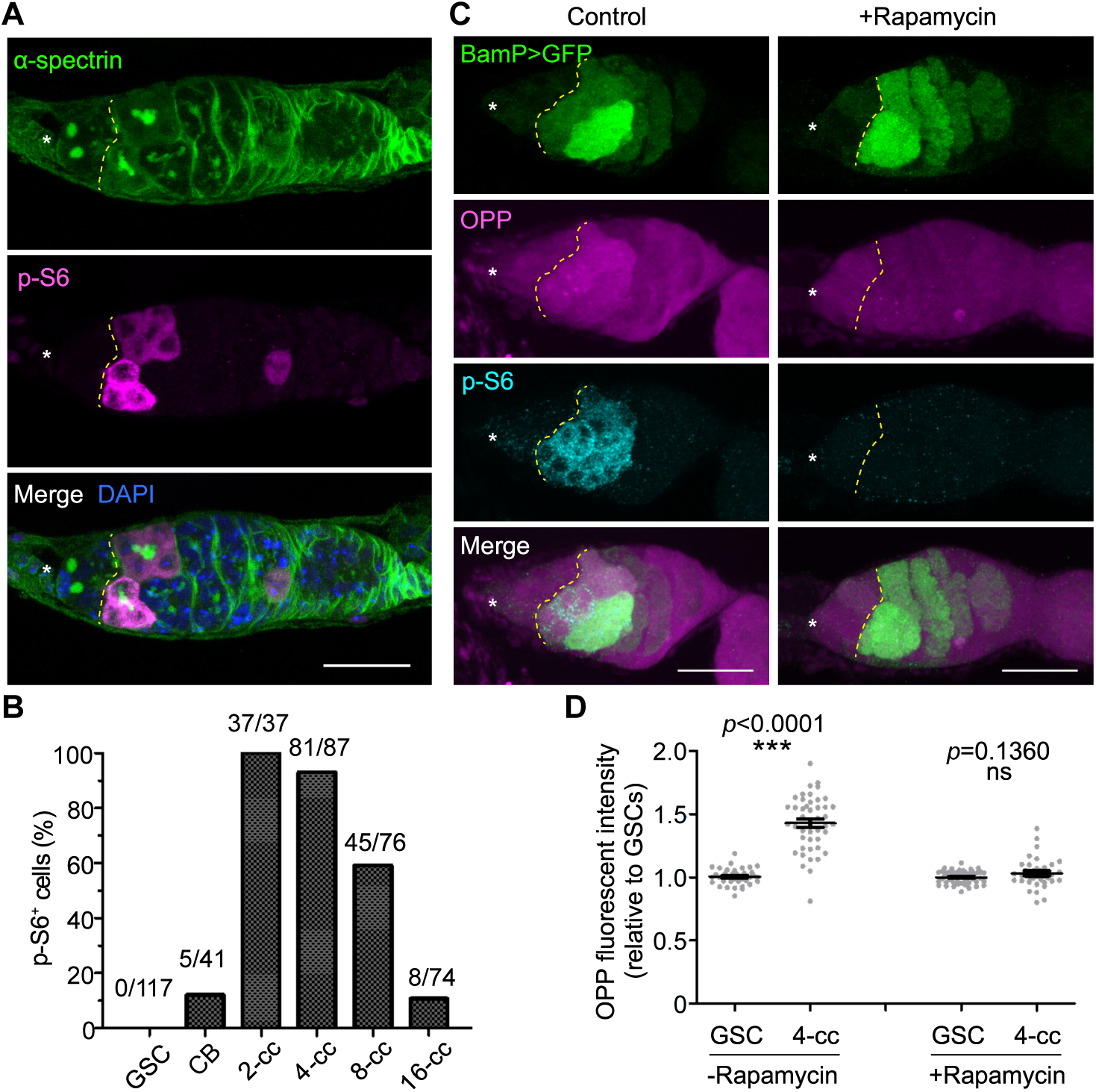
Tor is activated during germline stem cell differentiation and drives the observed increase in translation. (**A**) Representative image of a wild-type germarium labeled with α-spectrin (spectrosomes/fusomes, green), p-S6 (Tor activity, magenta), and DAPI (nuclei, blue). (**B**) Quantitation of the proportion of p-S6^+^ cells at different stages of germline differentiation. (**C**) Representative germaria with or without 10 μM rapamycin treatment, expressing the differentiation marker BamP>GFP (green), labeled with OPP (translation rate, magenta) and p-S6 (cyan). (**D**) OPP fluorescent intensity measurements of germaria with or without rapamycin treatment. Data are mean ± s.e.m \*\*\**p*<0.0001, t-test. Asterisks indicate the GSC niche. Dashed lines indicate the boundary between GSCs and differentiating cells (A, C). Scale bars, 20 μm (A, C).

Differentiating germ cells are characterized by a significant increase in protein synthesis rate (Sanchez et al., 2016), which temporally coincides with our observation of Tor activation. To test whether Tor mediates the increase in translation, we measured global protein synthesis rates *in vivo* using an imaging-based assay for O-propargyl-puromycin (OPP) incorporation into nascent polypeptides, which serves as a proxy for translation output (J. Liu et al., 2012; Sanchez et al., 2016). The robust increase in OPP incorporation observed in control differentiating cells was abolished by a short incubation with rapamycin (Figure 1C, D). Therefore, these results demonstrate that Tor is developmentally activated during cyst differentiation and plays a major role in promoting the observed increase in translation.

### Tor activation during GSC differentiation depends on the amino acid sensing pathway, not the insulin pathway

To determine which of the upstream molecular pathways participate in Tor kinase activation during GSC differentiation, we took advantage of tissue-specific RNAi knockdown of key pathway components (Figure 2A). As expected, p-S6 signal was abolished upon knockdown (KD) of either the Tor kinase itself or the downstream effector kinase ribosomal protein S6 kinase (S6K) (Figure 2B). Additionally, KD of Tsc1, a direct inhibitor of Tor kinase activity, resulted in a low, uniform p-S6 expression throughout the germarium, including the GSCs (Figure 2B).

**Figure 2.**
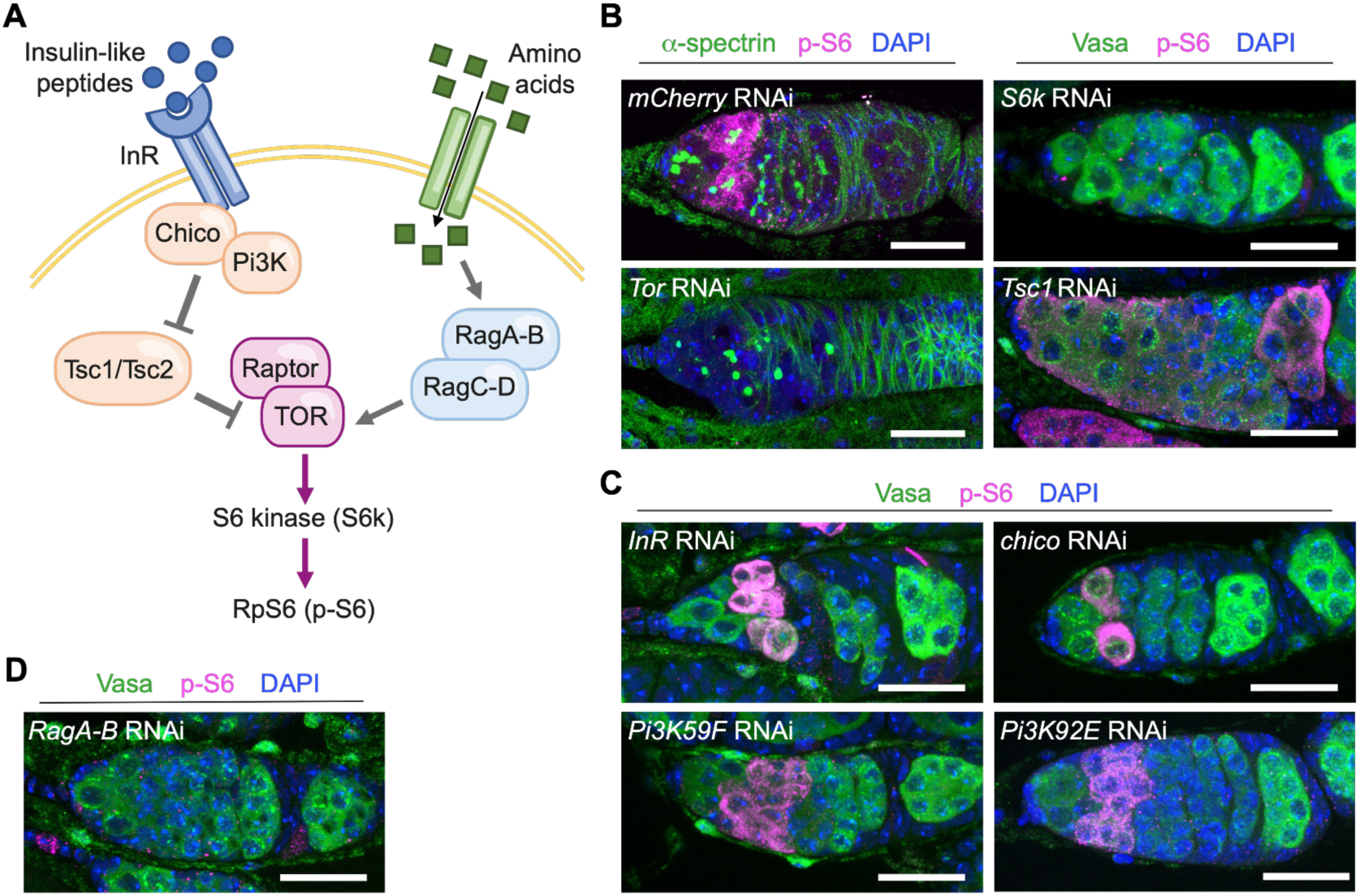
RagAB is required for Tor activation during GSC differentiation. (**A**) A simplified schematic showing insulin and amino-acid sensing pathways upstream of the Tor kinase, depicting the genes that were tested here. (**B**) Representative germaria of germline-specific KD of control (*mCherry* RNAi), *Tor, S6k* or *tsc1* labeled with α-spectrin (spectrosomes/fusomes, green) or Vasa (germline marker, green), p-S6 (Tor activity, magenta), and DAPI (nuclei, blue). (**C-D**) Representative germaria of germline-specific KD of *InR, chico, Pi3K59F (Vps34), Pi3K92E (Dp110)* or *RagAB* labeled with Vasa (green), p-S6 (magenta), and DAPI (blue). Scale bars, 20 μm (**B-D**).

The insulin receptor (InR)/phosphoinositide 3-kinase (PI3K)/AKT signaling cascade is one of the most established upstream activators of the Tor kinase during cell growth (Figure 2A). However, p-S6 expression was minimally affected upon KD of key components of the InR/PI3K cascade, including InR, chico/insulin receptor substrate (IRS), and the PI3K catalytic subunits Dp110/Pi3K92E and Vps34/Pi3K59F (Figure 2C). These data indicate that the InR/PI3K cascade is not a major activator of the Tor kinase during GSC differentiation, in agreement with the previous observation that the null *InR* mutant had a much less severe cyst growth delay than the null *Tor* mutant (LaFever et al., 2010). A second established upstream pathway regulating Tor depends on amino acid sensing, which can activate the Tor kinase through the Rag GTPases RagAB and RagCD (Figure 2A). When knocking down RagAB in germ cells, we observed that the p-S6 expression during GSC differentiation was abolished, similar to what was observed when knocking down Tor or S6K (Figure 2D). This finding suggests that the amino acid sensing module is upstream of Tor activation during GSC differentiation, rather than the InR/PI3K pathway.

### Mei-P26 activates Tor during GSC differentiation, uncoupling protein synthesis and ribosome biogenesis

Mei-P26, a TRIM-NHL protein ortholog of the mammalian TRIM32 family of proteins, has been reported to negatively regulate nucleolar size (a proxy for RiBi) during GSC differentiation (Neumüller et al., 2008). Phenotypically, *mei-P26*^*mfs1/mfs1*^ mutants initiate the GSC differentiation program, but differentiating cysts show enlarged nucleoli, increased cellular volume and are unable to complete terminal differentiation (Figure 3A,B) (Neumüller et al., 2008). Surprisingly, our analysis revealed that differentiating cysts in *mei-P26*^*mfs1/mfs1*^ mutants were also devoid of p-S6 signal, suggesting that Mei-P26 is required for Tor activation during differentiation (Figure 3B, C). Furthermore, the increase in global translation rate that is observed in differentiating cysts was abolished in *mei-P26*^*mfs1/ mfs1*^ mutants (Figure 3 D, E).

**Figure 3.**
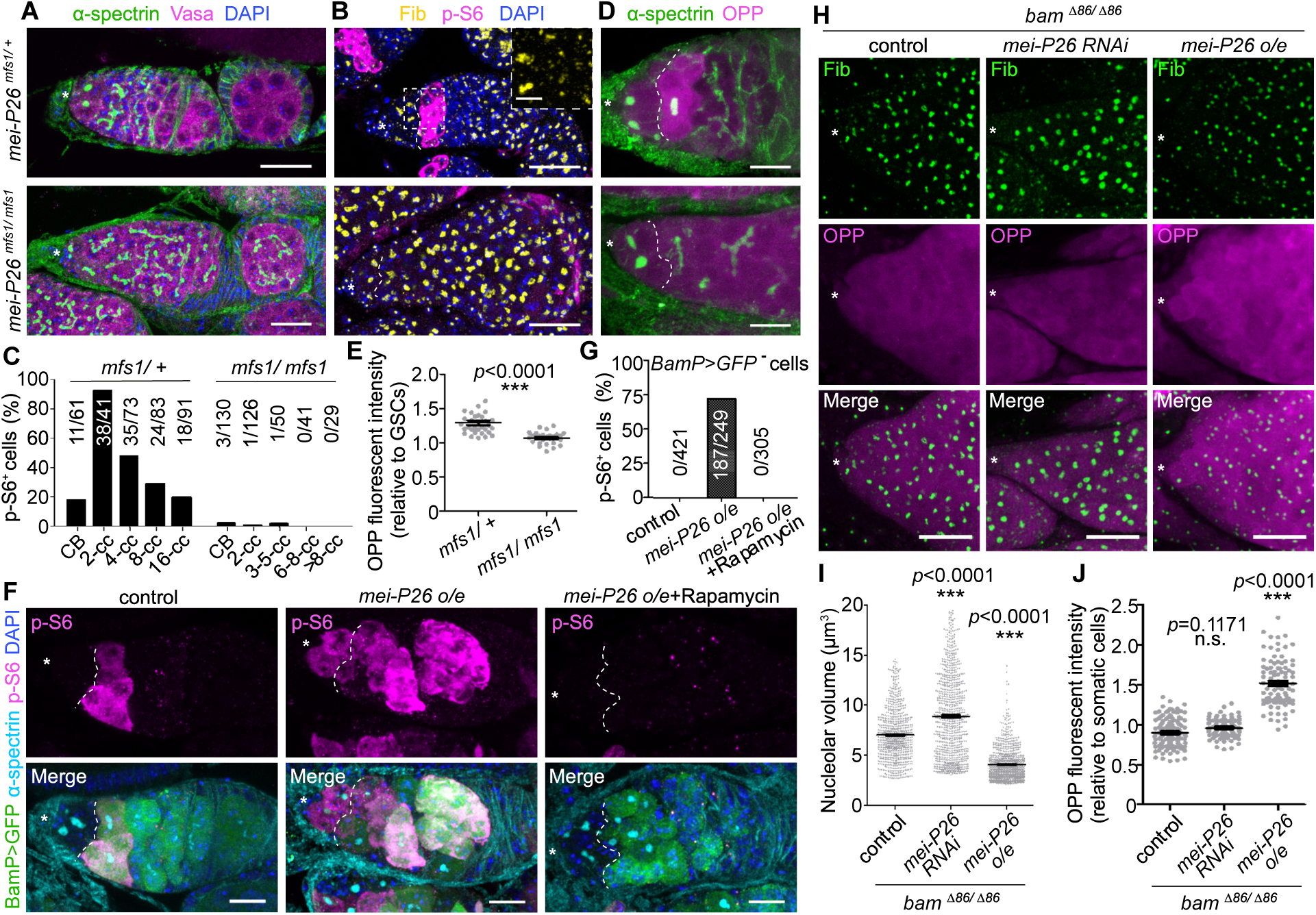
Mei-P26 activates Tor kinase during GSC differentiation. (**A, B** and **D**) Representative germaria of *mei-P26*^*mfs1/+*^ and *mei-P26*^*mfs1/mfs1*^ flies stained with α-spectrin (spectrosomes/fusomes, green, A and D), Vasa (germline, magenta, A), DAPI (nuclei, blue, A and B), Fib (nucleoli, yellow, B), p-S6 (Tor activity, magenta, B), and OPP (translation rate, magenta, D). (**C**) Quantitation of the proportion of p-S6^+^ cells at different stages of germline differentiation from *mei-P26*^*mfs1/+*^ and *mei-P26*^*mfs1/mfs1*^ flies. (**E**) OPP fluorescent intensity measurements of 2-4-cell cysts (cc) relative to GSCs, from germaria of *mei-P26*^*mfs1/+*^ or *mei-P26*^*mfs1/mfs1*^ flies. (**F**) Representative germaria of control (*nos-gal4/+*), and mei-P26 overexpression (o/e) (*nos-gal4/UASp-mei-P26)* flies with or without rapamycin feeding, stained with p-S6 (magenta), α-spectrin (cyan), GFP (BamP>GFP, differentiating cells, green) and DAPI (blue). (**G**) Proportion of p-S6^+^ GSCs in germaria of control (*nos-gal4/+*), and Mei-P26 o/e (*nos-gal4/UASp-mei-P26)* with or without rapamycin feeding. (**H**) Representative germaria of control (*nos-gal4/+*), *mei-P26* KD (*nos-gal4/UAS-mei-P26 RNAi)* or *mei-P26* o/e (*nos-gal4/UASp-mei-P26)* in the *bam*^*Δ86/Δ86*^ background, stained with Fib (green) and OPP (magenta). (**I** and **J**) Measurements of nucleolar volume (I) and translation rates (J) in GSC-like cells (*bam*^*Δ86*^) in germaria of *mei-P26* KD (*nos-gal4/UAS-mei-P26 RNAi)* or *mei-P26* o/e (*nos-gal4/UASp-mei-P26)* in the *bam*^*Δ86/Δ86*^ background. Data are mean ± s.e.m. \*\*\**p*<0.0001, t-test (E, I and J). Asterisks indicate the GSC niche. Dashed lines indicate the boundary between GSCs and differentiated cells (A, B, D and F). Scale bars, 20 μm (A, B, F and H), 10 μm (D), or 5 μm (B, inset).

To determine whether Tor kinase activity is directly promoted by Mei-P26, we overexpressed Mei-P26 in wild-type germ cells, which resulted in >75% (187 out of 249) of GSCs showing ectopic p-S6 signal (Figure 3F,G). In addition, extensive p-S6 signal was detected in fully differentiated egg chambers in *mei-P26* overexpressing (OE) ovaries (Supplementary figure 2A). It has been shown that sustained *mei-P26* OE induces premature GSC differentiation, eventually leading to loss of germ cells (Neumüller et al., 2008). We found that feeding flies with rapamycin not only abolished p-S6 signal in *mei-P26* OE ovaries but was also sufficient to suppress the GSC loss phenotype (Figure 3F, G, Supplementary figure 2A, B), indicating that ectopic Tor activation underlies Mei-P26-dependent premature loss of GSCs.

To characterize the molecular effects of modulating Mei-P26 expression in GSCs, we coupled overexpression and knockdown of Mei-P26 with a loss-of-function mutation for the differentiation factor Bag-of-marbles (Bam) (Mckearin & Spradling, 1990), which completely blocks GSC differentiation (Supplementary figure 2C). Immunofluorescence analyses revealed that OPP incorporation was positively correlated with Mei-P26 levels in *bam*^*Δ86*^ GSC-like cells while confirming that nucleolar volume displays an inverse correlation with Mei-P26 expression (Figure 3H-J) (Neumüller et al., 2008). Indeed, RNA-sequencing experiments on these *bam*^*Δ86*^ GSC-like cells with either *mei-P26* KD or OE revealed that RiBi-related genes – including those encoding ribosomal proteins, nucleolar markers, and ribosomal RNA (rRNA) polymerases – were negatively regulated by Mei-P26 (Supplementary figure 2D, Supplementary table 1). Our results demonstrate that Mei-P26 activates Tor to promote increased translation, and together with the previously shown role of Mei-P26 to suppress RiBi, this uncouples two of the most important anabolic processes during germline differentiation.

### Mei-P26 regulates RiBi through the pseudokinase TRRAP

It was previously shown that high levels of RiBi in the GSC are sustained mainly by TRRAP (Sanchez et al., 2016). TRRAP, encoded by Nipped-A in *Drosophila*, is the only pseudokinase in the phosphatidylinositol 3-kinase-related kinase (PIKK) protein family, which also includes Tor, ATM and ATR (Elías-Villalobos et al., 2019). While TRRAP is required for high RiBi in the GSC, its role during differentiation is unclear. To test whether the high rate of RiBi observed in differentiating *mei-P26* mutant cysts depends on TRRAP, we measured nucleolar size in *TRRAP* KD in the background of *mei-P26* KD. Enlarged nucleolar size in differentiating cysts in the *mei-P26* KD was significantly reduced upon additional *TRRAP* KD, compared to control *mCherry* KD (Figure 4A, B). As expected, KD of the Tor kinase only marginally affected the nucleolar size in the *mei-P26* KD (Figure 4A, B). This is in agreement with our observation that Tor activity is lost in the *mei-P26* KD, as well as the previous finding that Tor has a minimal role in the high RiBi rates observed in GSCs (Sanchez et al., 2016). Altogether, our data suggests that Mei-P26 regulates two PIKKs during differentiation, promoting translation through the activation of Tor while decreasing RiBi through TRRAP.

**Figure 4.**
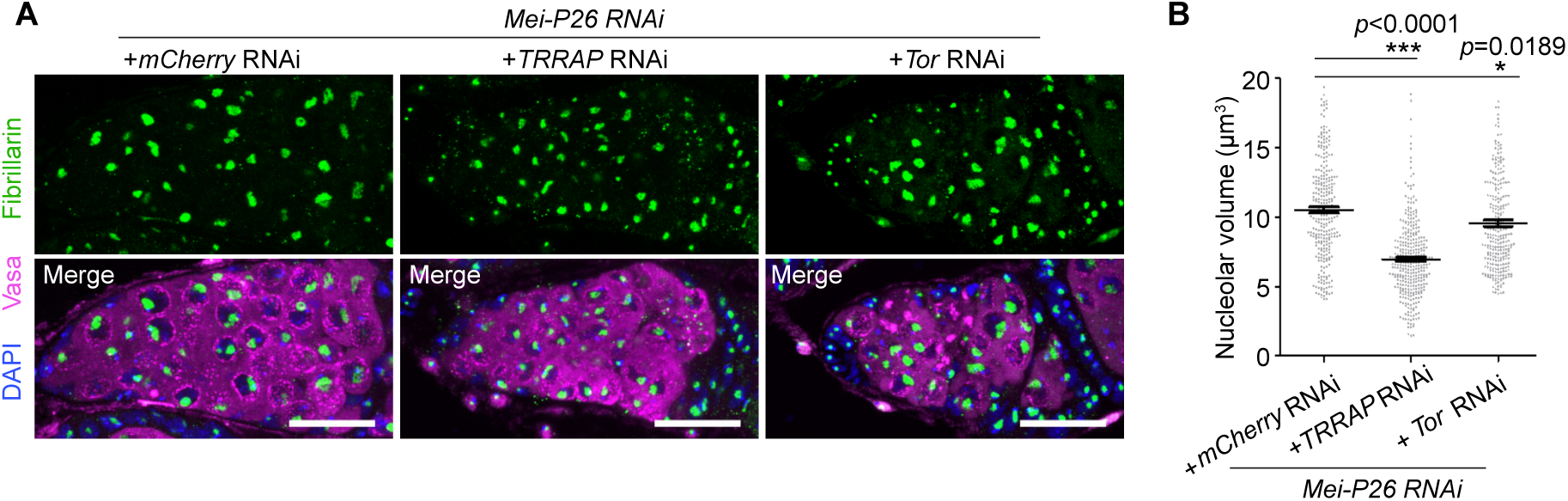
Mei-P26 suppression of RiBi depends on TRRAP/Nipped-A. (**A)** Representative germaria of germline-specific KD of *mei-P26* together with KD of *mCherry, TRRAP/ Nipped-A* or *Tor*, labeled with Fibrillarin (nucleolar marker, green), Vasa (germline marker, magenta), and DAPI (nuclei, blue). Scale bars, 20 μm. (**B**) Measurements of nucleolar volume in (A). Data are mean ± s.e.m. \*\*\**p*<0.0001, t-test (B).

### Redressing RiBi and Tor activities can suppress the defective differentiation and overgrowth phenotype induced by *mei-P26* mutants

Given that *mei-P26* mutants display defective differentiation, we investigated whether the misregulation of Tor activity and RiBi are responsible for the observed phenotypic changes. To do so, we modulated Tor and RiBi activities in *mei-P26-*depleted animals, aiming to restore the uncoupled metabolic environment that is characteristic of differentiation: high translation with low RiBi. As genetic manipulations of RiBi results in either a failure in differentiate or premature loss of GSCs (Sanchez et al., 2016), we took advantage of the small molecule RNA PolI inhibitor BMH-21, which has been shown to suppress rRNA synthesis in mammalian cells (Peltonen et al., 2014), to modulate RiBi. To test the effect of BMH-21 treatment on *Drosophila* ovaries, we used an assay based on the incorporation of 5-ethynyl uridine (EU) to image nascent RNA transcription with and without BMH-21 treatment. In controls, the bulk of the EU incorporation signal is generated by the Pol-I-mediated transcription of the ribosomal DNA repeats, accumulating in the nucleolus. Consistent with experiments in mammalian cells, we found that 100 μM BMH-21 treatment was sufficient to dramatically reduce transcription from the nucleolus in *Drosophila* ovaries (Supplementary figure 3).

We used BMH-21 treatment to test the effect of RiBi suppression on *mei-P26*-depleted germ cells. We observed that abdominal injection of BMH-21 into adult flies led to a moderate suppression of the *mei-P26* KD-induced block in germline differentiation, with ∼18% of ovarioles (*mei-P26* RNAi, *mCherry* RNAi + BMH-21) containing terminally differentiated egg chambers (Figure 5A, B). A similar suppression effect was observed by genetically activating the Tor kinase during germline development by knocking down Tsc1 or Tsc2 (*mei-P26* RNAi, *tsc1* RNAi or *tsc2* RNAi), resulting in 12-15% of ovarioles with terminally differentiated egg chambers (Figure 5A, B). When RiBi was suppressed at the same time as activating Tor, robust terminal differentiation was observed in ∼40% of ovarioles, and ∼10% of ovarioles contained fully developed eggs (Figure 5A, B). These experiments show that the block in differentiation phenotype observed in the *meiP26* mutant can be significantly suppressed by redressing levels of Tor activation and RiBi, suggesting that the uncoupled state is essential for proper differentiation.

**Figure 5.**
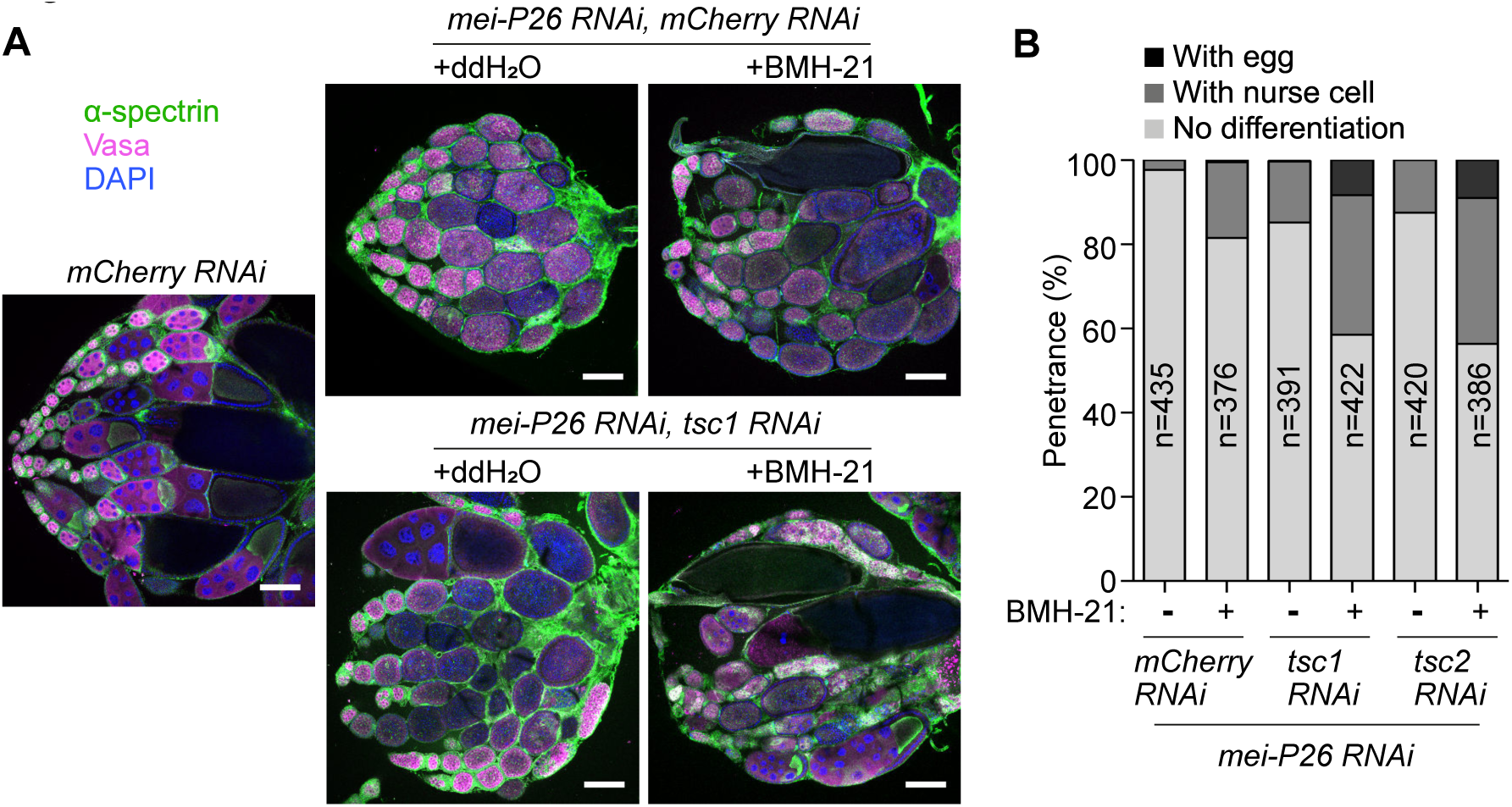
Manipulating RiBi and Tor activity in *mei-P26* ovaries can promote terminal differentiation. **(A)** Representative ovaries of control (*mCherry RNAi*), *mei-P26 mCherry* double KD (*mei-P26 RNAi, mCherry RNAi*), and *mei-P26 tsc1* double KD (*brat RNAi, tsc1 RNAi*) flies, with or without BMH-21 feeding. Samples were stained with α-spectrin (spectrosome/fusomes, green), Vasa (germline, magenta), and DAPI (nuclei, blue). **(B)** Penetrance of ovary phenotypes of control (*mCherry RNAi*), *mei-P26 mCherry* double KD (*mei-P26 RNAi, mCherry RNAi*), *mei-P26 tsc1* double KD (*mei-P26 RNAi, tsc1 RNAi*), and *mei-P26 tsc2* double KD (*mei-P26 RNAi, tsc1 RNAi*) with or without BMH-21 injection. *n* is the number of ovarioles analyzed. Scale bars, 200 μm (A).

### Brat is required for Tor activation during neuroblast differentiation

To determine whether TRIM-NHL proteins regulate Tor activity in other stem cell differentiation systems, we examined the type II NBs during larval neurogenesis. In this system, the Mei-P26 paralog, Brain tumor (Brat), also promotes differentiation while downregulating RiBi activity (Betschinger et al., 2006; Frank et al., 2002). In the developing larval brain, Brat activity is restricted to differentiating progeny of the type II NBs by protein segregation during an asymmetric cell division (Betschinger et al., 2006; Lee et al., 2006). As in the case of GSCs, NBs are characterized by a larger cell size and a higher RiBi rate compared to their differentiating progeny, which include immature intermediate neural progenitors (INPs), mature INPs, ganglion mother cells (GMCs) and neurons (Supplementary figure 4A) (Betschinger et al., 2006; Homem & Knoblich, 2012). First, we examined Tor activation during type II NB differentiation using immunofluorescence to detect p-S6. We detected p-S6 in ∼64% of type II NBs (large cells; Deadpan, Dpn^+^), ∼73% of their immediate progeny cells (immature INPs: Dpn^-^; Prospero, Pros^-^) and ∼80% of mature INPs (Dpn^+^, Pros^-^). However, fewer than 11% of differentiated neurons (Pros^+^) were positive for p-S6 (Figure 6A, B). We examined the effect of rapamycin treatment on pS6 in the type II lineage. We found that 20 minutes of rapamycin treatment during *ex vivo* brain culture led to substantial loss of p-S6 signal in the NBs and INPs (Supplementary figure 4B). Therefore, we conclude that Tor activity is developmentally regulated during type II NB differentiation.

**Figure 6.**
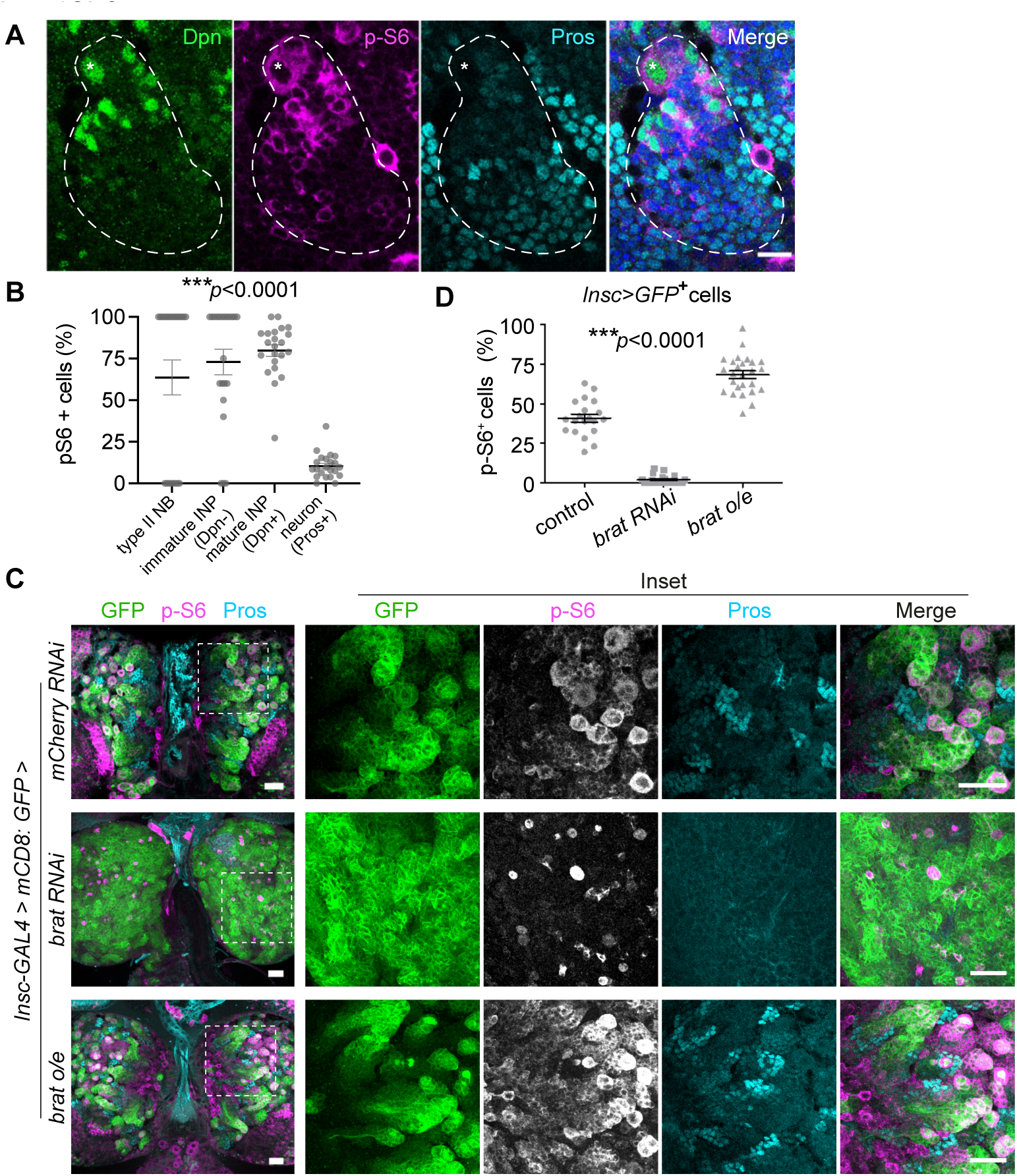
Brat is required for Tor activation during type II NB differentiation. (**A**) Representative confocal image of a type II NB lineage in the L3 larval brain, stained with Dpn (self-renewal marker, green), p-S6 (Tor activity, magenta), and Pros (pro-differentiation marker, cyan). Dashed line surrounds a type II NB lineage, asterisk indicates a type II NB. (**B**) Proportion of p-S6^+^ cells during type II NB differentiation. Each point is a single cluster analyzed, and each cluster contains a single type II NB. (**C**) Representative larval brains of control (*mCherry* RNAi), *brat* RNAi or *brat* overexpression (o/e) driven by *insc-GAL4*, stained with GFP (*insc>mCD8:GFP*, green), p-S6 (magenta or gray) and Pros (cyan). White dashed rectangles (left) indicate the sources of the insets (right). (**D**) The percentage of p-S6^+^ GFP^+^ in control (*mCherry RNAi*), *brat* RNAi, and *brat* o/e larval brains. Data are mean ± s.e.m. \*\*\**p*<0.0001, one-way ANOVA (B, D). Scale bars, 10 μm (A) or 25 μm (C).

We tested whether Brat is required for Tor activation during NB differentiation, using *brat* RNAi driven by *insc*-Gal4, which drives in NBs and INPs. Brat functions in the immature INPs to promote INP maturation and prevent reversion to a NB fate (Xiao et al., 2012), therefore *brat* KD leads to an accumulation of supernumerary NBs. Although the *brat* KD brain was filled with NB-like cells, we found that that p-S6 signal was mostly abolished in *brat* deficient brains (Figure 6C, D), becoming restricted to a small number of cells, whose number and distribution resembled that of NBs in wild-type brains. This result suggests that while NBs sustain Tor activation in a Brat-independent manner, upon NB division, Tor activity in progeny cells relies on Brat. To test whether Brat is sufficient to promote Tor activation, we investigated the expression of p-S6 in larval brains with Brat overexpression. In this context, the p-S6 expression domain was robustly and ectopically expanded into GMCs and newly differentiated neurons (Figure 6C, D). These results demonstrate that, analogous to the role of Mei-P26 in GSC differentiation, Brat activates the Tor kinase alongside its known role of downregulating RiBi during differentiation (Betschinger et al., 2006; Frank et al., 2002).

### The phenotype of *brat*-deficiency in larval brains can be suppressed by redressing RiBi and Tor activity

We showed that the phenotype of *mei-P26* mutants in the ovary can be rescued by simultaneously suppressing RiBi and activating Tor, to restore the metabolic uncoupling that is characteristic of differentiation. Therefore, we investigated whether modulating Tor and RiBi activities in the larval brain could also circumvent the block to differentiation observed in *brat-*depleted brains. *brat* depletion leads to the accumulation of undifferentiated Dpn^+^ Pros^-^ cells. Clonal analyses using the *brat*^*11*^ mutation revealed that knockdown of the Tor-antagonizing regulators Tsc1 or Tsc2 (Figure 2A) was sufficient to restore differentiation (Dpn^-^ Pros^+^ cells) in the absence of Brat in about 10% of clones (2 in 15 clones for *tsc1* RNAi or 3 in 31 clones for *tsc2* RNAi, Supplementary figure 5A).

Moreover, we found that increasing Tor activity through overexpression of Raptor (Figure 2A) in a *brat* RNAi context (*brat* RNAi, *UAS-raptor:HA*) was sufficient to decrease the undifferentiated Dpn^+^ Pros^-^ population and to promote differentiation (Dpn^-^ Pros^+^ cells; Figure 7). Indeed, in *brat* RNAi larvae with Raptor OE, ∼50% of brain lobes had no apparent phenotypic abnormalities (*mCD8:GFP*; Figure 7, Supplementary figure 5B,C). This cellular rescue resulted in a functional recovery: climbing assays revealed that while *brat*-deficient flies had reduced mobility, adult flies overexpressing Raptor in a *brat* RNAi context were indistinguishable from control flies (Supplementary figure 6A,B, Supplementary data video 1). These results show that Tor activation can promote the differentiation of *brat*-deficient NBs.

**Figure 7.**
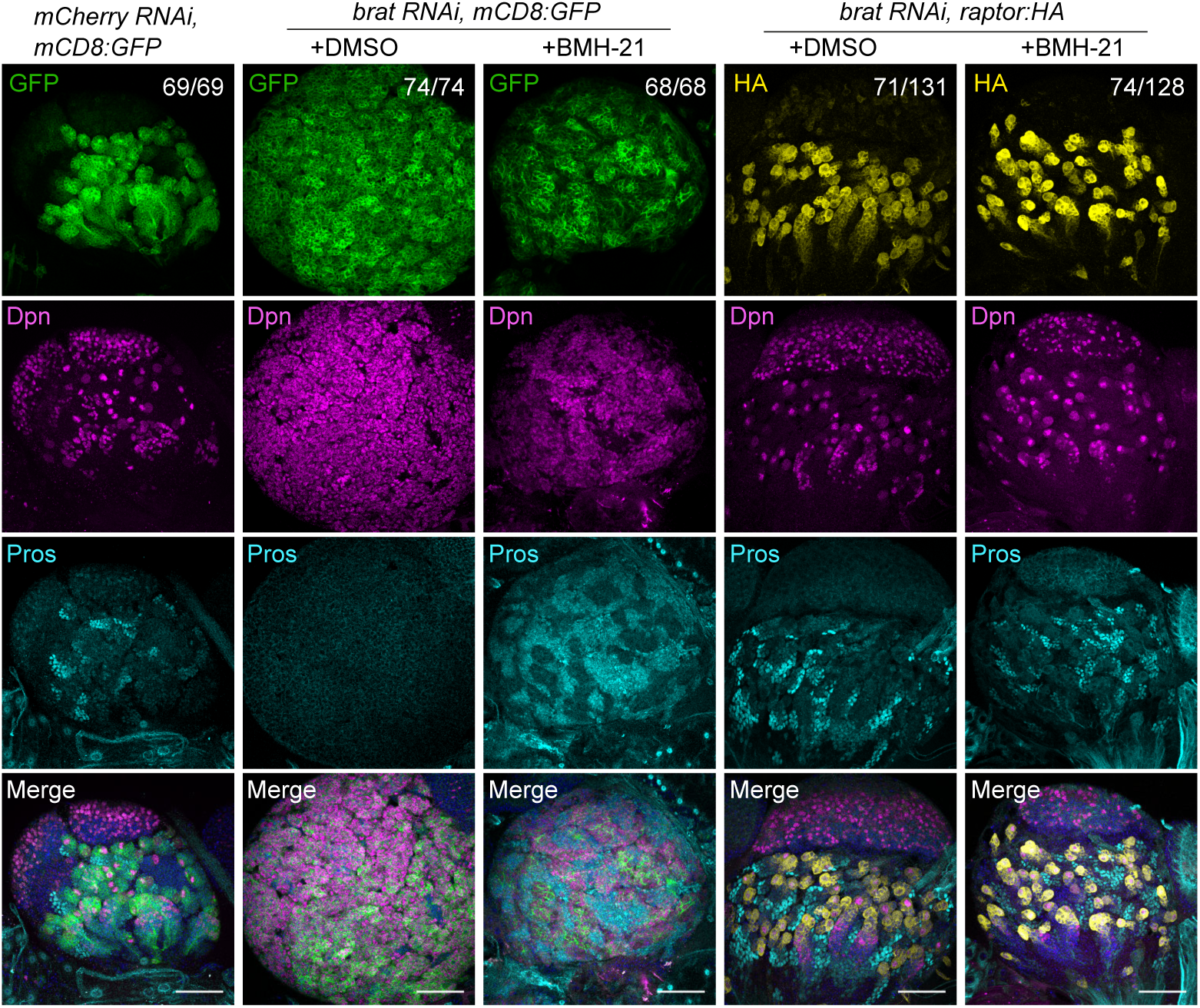
Redressing RiBi and Tor activity promotes terminal differentiation in *brat* larval brains. (**A**) Representative larval brain lobes of control (*mCherry RNAi, mCD8:*GFP), *brat* KD (*brat RNAi, mCD8:*GFP), and *brat* KD with *raptor* overexpression (*brat RNAi, raptor:HA*) flies (driven by *insc-GAL4*) with or without BMH-21 feeding. Samples were stained with GFP (mCD8, green) or HA (Raptor, yellow), Dpn (self-renewal marker, magenta), Pros (pro-differentiation marker, cyan), and DAPI (nuclei, blue). Numbers in the top panel indicate the penetrance of phenotypes out of three independent experiments. Scale bars, 50 μm.

To determine whether inhibition of RiBi has a similar effect on the differentiation of *brat*-deficient NBs, we fed larvae with BMH-21. *brat* RNAi brains showed partial differentiation after BMH-21 feeding, with increased expression of the differentiation marker Pros (Figure 7, Supplementary figure 7A, B). However, the distribution of cellular markers did not fully recapitulate the wild-type pattern and we observed numerous cells positive for both Pros and Dpn markers, suggesting that the canonical program of type II NB differentiation was not fully restored (Supplementary figure 7B, yellow arrows). To examine the effect of genetic RiBi manipulation, we knocked down Nop60B, a pseudouridine synthase involved in rRNA processing. Remarkably, *Nop60B* KD suppressed the accumulation of Dpn^+^ Pros^-^ NB-like cells observed in *brat* KD and restored the differentiation pathway to produce Dpn^-^ Pros^+^ cells (Supplementary figure 7C).

Collectively, our results demonstrate that TRIM-NHL proteins drive terminal differentiation by simultaneously suppressing RiBi and promoting translation, uncoupling two of the most energy-consuming biosynthetic activities.

## Discussion

Accumulating evidence indicates that dynamic regulation of cellular metabolism plays a key role at the tissue level in development and homeostasis (Buszczak et al., 2014; Miyazawa & Aulehla, 2018). During stem cell differentiation, RiBi and protein synthesis rates are actively regulated, and these dynamic changes are essential for the balance of self-renewal, growth, and differentiation (Teixeira & Lehmann, 2019). Here, we have shown that two members of the TRIM-NHL protein family (Mei-P26 and Brat) are responsible for the simultaneous regulation of RiBi and Tor kinase activity/translation during stem cell differentiation, in GSC and NB systems. The antagonistic regulation of these highly energy-consuming processes creates a state of high translation with low RiBi, which deviates from the canonical growth paradigm downstream of Tor activation (G. Y. Liu & Sabatini, 2020). We find that the differentiation block in *mei-P26/brat* mutants can be rescued by modulating translation and RiBi to restore the uncoupled state, suggesting that metabolic uncoupling is a driver of terminal differentiation.

We have shown that Mei-P26 and Brat uncouple translation and RiBi specifically during differentiation. However, both Mei-P26 and Brat are expressed in the stem cells as well as their differentiating progeny (Lee et al., 2006; Neumüller et al., 2008), so it remains an open question how their activity is restricted to the differentiating cells. Brat is transcribed and translated in NBs, but is sequestered to the basal membrane and segregated into differentiating daughter cells (Lee et al., 2006). This mechanism enriches Brat in the INP compared to the NB and perhaps this higher concentration is necessary for Brat action. In contrast, Mei-P26 is equally expressed between GSCs and differentiating cells (Neumüller et al., 2008), suggesting that a different regulatory mechanism is at play in the GSC to limit Mei-P26 activity. Interestingly, we have shown that overexpression of Mei-P26 is sufficient to drive ectopic Tor activation in the GSC, suggesting that the as yet unknown factor that suppresses Mei-P26 activity in GSCs becomes overwhelmed in a Mei-P26 overexpression condition.

In any case, the presence of Mei-P26 protein from the GSC stage may allow for the prompt modulation of its downstream targets in the first stages of differentiation. Indeed, our results indicate that Mei-P26 acts very rapidly in the early stages of differentiation, regulating two different PIKK members in opposite directions: promoting translation through activating Tor, and suppressing RiBi through TRRAP. Mei-P26 upregulates the activity of the Tor kinase but the molecular mechanisms of its interaction with TRRAP are unknown. Interestingly Tor is activated in cystoblasts, but nucleolar volume reduction is only apparent from the 2-cell cyst stage onwards (Sanchez et al., 2016), which may be explained by the rapid action of the Tor kinase compared to TRRAP, which is a transcriptional regulator (Elías-Villalobos et al., 2019).

Contrasting the functions and protein structures of Mei-P26 and Brat can provide some insight into the potential mechanisms of action at the molecular level. TRIM-NHL family members are generally characterized by an N-terminal TRIM motif, which can mediate protein-protein interactions and enable ubiquitin-mediated protein degradation, and a C-terminal NHL domain, which can mediate RNA binding (Tocchini & Ciosk, 2015). Notably, Brat lacks a RING domain (Loedige et al., 2014, 2015)and yet activates Tor and suppresses RiBi, suggesting that the RING domain is unlikely to be necessary for these activities, unless Brat and Mei-P26 act through different mechanisms. Furthermore, Brat is also expressed in the ovary, in differentiating GSC progeny where it has a role in cell size and differentiation (Harris et al., 2011). It is unclear how Mei-P26 and Brat may interact in the ovary and it will be important to build a more complete molecular picture of Mei-P26 and Brat activity.

Unlike in GSCs, Tor is active in NBs. Upon *brat* KD, INPs revert to a NB-like fate (Xiao et al., 2012), but we observed that Tor activity was not maintained, except in a small number of cells resembling the original NBs. Given that Brat is asymmetrically segregated into the differentiating daughter cells, it would not be surprising if Tor activity in the type II NBs is Brat-independent. Notably after type II NB division, Tor activity in progeny cells becomes dependent on Brat, which implies the existence of a mechanistically complex switch that is required to maintain Tor activation from NBs through the first stages of differentiation. In late stage larvae, Tor activation in the type I NB is independent of insulin or amino acid signaling, and is instead activated by ligands expressed from the glial niche (Cheng et al., 2011). In this so-called brain-sparing mechanism, glial signaling secures NB growth and division regardless of nutritional status, ensuring the successful production of the complete brain during larval development. In sharp contrast to the brain-sparing process, GSC growth and proliferation is acutely dependent on the nutritional state of the animal, in order to maximize survival of the offspring (Drummond-Barbosa & Spradling, 2001). Therefore it is not surprising that we find a role for amino-acid sensing in the activation of Tor kinase during GSC differentiation.

Tor kinase activity is generally associated with growth (G. Y. Liu & Sabatini, 2020), but interestingly in both GSC and NB systems, differentiation is accompanied by a decrease in cellular volume while Tor is active (Homem et al., 2013; Neumüller et al., 2008). Furthermore, *mei-P26* and *brat* mutants display tumor phenotypes of large, undifferentiated cells despite the loss of Tor activity (Betschinger et al., 2006; Neumüller et al., 2008). Canonically, Tor activation is associated with an increase in both translation and ribosome biogenesis, creating an optimal scenario to drive growth (G. Y. Liu & Sabatini, 2020). However, in the stem cell systems described here, Tor activity is uncoupled from ribosome biogenesis such that limited ribosomes are produced even while Tor is active. This lack of ribosome generation may act as a brake on Tor-driven growth. We must also consider that the observed changes in cell size are likely to be affected by both cell growth and division. Tor has been shown to promote germline cyst proliferation (LaFever et al., 2010). In the *mei-P26*/*brat* tumor scenario, the loss of Tor may slow cell divisions such that cells have more time to grow, while the increased RiBi rate might allow growth to continue beyond its usual depletion of resources. This combination would result in larger cells despite a lower growth rate.

In this way, the relationship between growth and cell cycle may also provide insight into how uncoupling of translation and RiBi drives differentiation. Cell cycle exit is a prerequisite for terminal differentiation in both the GSC and NB systems (Hinnant et al., 2020; Homem & Knoblich, 2012). In general, the coordination of anabolic activities is required for sustained growth and proliferation (Bracha Ginzberg et al., 2018), so perhaps uncoupling translation and Ribi limits the possible growth and number of cell divisions prior to exhaustion of cellular resources. In the pupal brain, type I NB growth is limited such that cells shrink with each division and this shrinkage is required to induce the final symmetric division that leads to terminal differentiation of both daughters (Homem et al., 2014; Yang et al., 2017). In the GSC and larval NB systems studied here, anabolic uncoupling may start a timer before resource depletion leads to terminal differentiation. Careful analysis of proliferation rates and cell size, perhaps using long term live imaging, will be needed to unpick the contributions of growth and division.

Altogether, our work uncovers a hitherto overlooked role of metabolic imbalance in driving differentiation. As RiBi and translation rates are known to be dynamically regulated in many different stem cell systems, we propose that this mechanism has general implications for our understanding of differentiation.

## Materials and Methods

### *Drosophila* stocks, genetics, and husbandry

*Drosophila melanogaster* stocks and transgenes used: *w*^*1118*^ (Lehmann lab stock); *UAS-Dcr2, w*^*1118*^; *nosP-GAL4-NGT40* (BDSC #25751); *P{bamP-GFP}*transgene (Chen & McKearin, 2003); *bam*^*Δ86*^, *ry, e/TM3, Sb, ry, e* (McKearin & Ohlstein, 1995); *y*^*1*^ *w*^*1*^ *mei-P26*^*mfs1*^;;; *Dp(1;4)A17/sv*^*spa-pol*^ (BDSC #25919) (Page et al., 2000); *w*^*1118*^; *UASp-mei-p26*.*N* (BDSC #25771); *brat*^*11*^; *insc-Gal4 (BDSC #8751)* and UAS*-mCD8:GFP* (BDSC #5130 and #5137); UASp-*brat (Harris et al*., *2011)*; UAS-*raptor-HA* (BDSC #53726); *w*; FRT 40A* (BDSC #86317); *hs*-*FLP, UAS-mCD8: GFP; tubP-GAL80, FRT 40A; tubP-GAL4* (from BDSC #44406 and #84300); *w;; [FlyFos020668(Tor29074::2XTY1-SGFP-V5-preTEV-BLRP-3XFLAG)dFRT] VK00033* (VDRC #318201), *w;; [FlyFos022619(raptor[16724]::2XTY1-SGFP-V5-preTEV-BLRP-3XFLAG)dFRT]VK00033* (VDRC #318149).

The following *UAS-RNAi* lines were used: *y*^*1*^ *sc*^*\**^ *v*^*1*^ *sev*^*21*^; *P{VALIUM20-mCherry}attP2* (*mCherry RNAi*; BDSC #35785); *y*^*1*^ *sc*^*\**^ *v*^*1*^; *P{TRiP*.*GL01124}attP40* (mei-P26-shRNA; BDSC #36855); *y*^*1*^ *sc*^*\**^ *v*^*1*^; *P{TRiP*.*HMS01121}attP2* (*brat RNAi*, BDSC #34646); *y*^*1*^ *sc*^*\**^ *v*^*1*^; *P{TRiP*.*GL00012}attP2* (*tsc1 RNAi*, BDSC #35144); *y*^*1*^ *sc*^*\**^ *v*^*1*^; *P{TRiP*.*GL00321}attP2* (*tsc2 RNAi*, BDSC #35401); *y[1] sc[*] v[1] sev[21]; P{y[+t7*.*7] v[+t1*.*8]=TRiP*.*GL00139}attP2* (*InR* RNAi, BDSC#35251); *y[1] sc[*] v[1] sev[21]; P{y[+t7*.*7] v[+t1*.*8]=TRiP*.*GL00139}attP2* (*chico* RNAi, BDSC#28329); *y[1] v[1]; P{y[+t7*.*7] v[+t1*.*8]=TRiP*.*HMS00007}attP2* (*Akt* RNAi, BDSC#33615); *y[1] sc[*] v[1] sev[21]; P{y[+t7*.*7] v[+t1*.*8]=TRiP*.*HMS01064}attP2* (*RagAB* RNAi, BDSC#34590); y*[1] sc[*] v[1] sev[21]; P{y[+t7*.*7] v[+t1*.*8]=TRiP*.*GL01327}attP2* (*s6k* RNAi, BDSC#41895); *y[1] sc[*] v[1];; P{TRiP*.*HMS00904}attP2 [Tor]* (*Tor* RNAi, BDSC#33951*); y[1] sc[*] v[1] sev[21]; P{y[+t7*.*7] v[+t1*.*8]=TRiP*.*HMC05152}attP40 (Pi3K92E/*Dp110 RNAi BDSC#61182); *y[1] sc[*] v[1] sev[21]; P{y[+t7*.*7] v[+t1*.*8]=TRiP*.*HMS00261}attP2/TM3, Sb[1]* (*Pi3K59F/Vps34* RNAi BDSC#33385); *y[1] sc[*] v[1] sev[21]; P{y[+t7*.*7] v[+t1*.*8]=TRiP*.*GL00156}attP2* (*LexA* RNAi, BDSC#67945); *y[1] sc[*] v[1] sev[21]; P{y[+t7*.*7] v[+t1*.*8]=TRiP*.*HMC04815}attP40* (*nop60B* RNAi, BDSC#57500), ;;*PBac{fTRG00888*.*sfGFP-TVPTBF}VK00033* (Nop60B::GFP, VDRC #318245)

Unless stated otherwise, stocks and crosses were maintained on standard propionic food at 25°C. For rapamycin feeding experiments with adult flies, 200 μl 100 μM rapamycin (Sigma Aldrich, #R0395) was added to the top of food at least one day before newly eclosed flies were transferred into the vial. Flies were raised in food containing rapamycin for 3 days at 25°C before ovary dissections. For feeding experiments with larvae, 50 μl 2 mM BMH-21 (Sigma Aldrich, #SML1183) was added to the top of food for no less than one day before the experiment. Three-day-old larvae grown at room temperature were transferred to the food containing BMH-21 and raised at 29°C for three days before brain dissections. For BMH-21 injection in adult flies, 69 nl 200 μM BMH-21 diluted in double-distilled water was injected into the abdomen of one-day-old females. Injected flies were raised for five days at 25°C before ovary dissections.

### Immunofluorescence and antibodies

Adult ovaries were dissected in cold PBS buffer and fixed in PBST (PBS with 0.2% Triton-X100) containing 4% Formaldehyde (Thermo Fisher Scientific, #28908) for 30 minutes. Larval brains were dissected in Schneider’s insect medium and transferred immediately into cold fixative (4% formaldehyde in PBST), then fixed for 25 minutes. Fixed tissues were rinsed three times with PBST before incubation in blocking buffer (PBS with 5% goat serum; Sigma Aldrich, #G9023) overnight at 4°C. Samples were then incubated with primary antibody diluted in blocking buffer overnight at 4°C, washed four times with blocking buffer, and incubated with secondary antibodies and DAPI (Sigma Aldrich, #D9542) diluted in blocking buffer overnight at 4°C. Samples were washed four times with PBST and mounted in VectaShield medium (Vector Laboratories, #H1000). Fluorescent images were acquired on a Leica SP8 confocal microscope using a 40X oil objective or a 20X dry objective. Images were processed using ImageJ (NIH; http://imagej.nih.gov/ij/).

The following antibodies were used for immunofluorescence staining: rat anti-Dpn (Abcam, #ab195173, 1:200), mouse anti-Fib (Abcam, #ab4566, 1:200), rabbit anti-HA (Abcam, #ab9110, 1:100), rat anti-HA (Sigma Aldrich, #3f10, 1:100), rat anti-GFP (Millipore, #MAB3580, 1:200), chicken anti-GFP (1:1000), rat anti-RFP (Chromotek, #5F8, 1:200), mouse anti-α-Spectrin (DSHB, #3A9, 1:100), mouse anti-Pros (DSHB, #MR1A, 1:20), rabbit anti-phosphorylated-S6 (Dr Aurelio Teleman, 1:200; and custom made antibody generated by Davids Biotechnologie, Germany, as previously described in (Romero-Pozuelo et al., 2017)), rabbit anti-Vasa (Dr Ruth Lehmann, 1:5000), rabbit anti-Mei-P26 (Dr. Paul Lasko, 1:1000), goat anti-mouse Alexa488 (Invitrogen, #A32723, 1:200), donkey anti-rat Alexa488 (Invitrogen, #A11006, 1:200), goat anti-rabbit Alexa568 (Invitrogen, #A11011, 1:200), donkey anti-mouse Cy3 (Jackson ImmunoResearch, #715-165-150, 1:200), donkey anti-rabbit Alexa647 (Jackson ImmunoResearch, #711-605-152, 1:200), and goat anti-mouse Alexa647 (Invitrogen, #A21236, 1:200).

### Measurement of global protein synthesis *in vivo*

Protein synthesis was detected by the Click-iT® Plus OPP Alexa Fluor 594 Protein Synthesis Assay Kit (Molecular Probes) as previously described (Sanchez et al., 2016). Unless stated otherwise, ovaries were dissected in Shields and Sang M3 Insect Medium (S3652, Merck) and were immediately transferred after dissection to fresh medium containing a 1:400 dilution of Click-iT OPP Reagent (OP-puro 50 μM). For the experiment in Figure 1C, dissected ovaries were incubated in media containing 10 μM rapamycin for 30 min before exposure to the Click-iT OPP reagent. Samples were incubated with OPP at room temperature for 30 min, rinsed 3 times with PBS, and fixed with 4% formaldehyde in PBS for 30 min. After the Click-iT reaction, samples were washed with PBS with 1% BSA and 0.2% Triton X-100 for 1 hour and immunostained according to standard procedures. Quantification of OP-Puro fluorescence intensity was performed as previously described (Sanchez et al., 2016) using ImageJ. Each experiment was performed at least three times.

### Measurement of nascent RNA synthesis with BMH-21 treatment

Ovaries were dissected in Shields and Sang M3 Insect Medium (S3652, Merck). Ovaries were treated for 1 hour at RT with 100 μM BMH-21 or DMSO control, in Shields and Sang M3 Insect Medium. We estimate 100 μM BMH-21 to be approximately equivalent to the concentration in the abdomen after injection in our experiments. Media was exchanged to also include 10 mM 5-ethynyl uridine (EU) and ovaries were incubated at room temperature for a further 2 hrs. Ovaries were fixed with 4% formaldehyde in PBS containing 0.3% Triton-X, for 25 min at RT, and then washed 3 times for 15 minutes each in PBS with 0.3% Triton-X. Nascent RNA was visualized using Click-iT RNA Alexa Fluor 594 Imaging Kit (C10330, ThermoFisher Scientific) according to manufacturer’s instructions.

### *ex vivo* brain culture

L3 larval brains were dissected in culture media (80% Schneider’s Insect Medium, 20% Fetal Bovine Serum and larval extract (according to (Hailstone et al., 2020) but with the omission of insulin). After dissection, brains were incubated at for 20 min room temperature in culture media with or without 10 μM rapamycin. Brains were then fixed and stained according to standard procedure described above.

### RNA sequencing

120 pairs of ovaries were dissected for each sample and immediately stored at −80°C after dissection. Frozen samples were homogenized in TRIzol Reagent (Invitrogen) using an electrical pestle and further disrupted by passing 15 times through a 26-gauge syringe. Total RNA was isolated using TRIzol Reagent (Invitrogen) following the manufacturer’s protocol. After RNA quantification using Qubit (Invitrogen), Poly(A)-selected RNA-sequencing (RNA-seq) libraries were generated using 2.5 μg of purified RNA with the NEBNext^®^ Poly(A) mRNA Magnetic Isolation Module and the NEBNext^®^ Ultra™ Directional RNA Library Prep Kit for Illumina^®^. Libraries were multiplexed using the NEBNext^®^ Multiplex Oligos for Illumina^®^ and sequenced in single-end, 50-nt-long reads on an Illumina HiSeq 2500. The resulting RNA-sequencing data was first aligned to ribosomal RNA using Bowtie2 (Langmead & Salzberg, 2012). Non-rRNA reads were mapped to the *Drosophila melanogaster* genome (dm6) using STAR (Dobin et al., 2013), and transcript abundance was quantified and differentially expressed genes were identified using Cufflinks (Trapnell et al., 2010). Analyses were performed with two samples, each with two biological replicates.

### Climbing assay

20 1-2 day-old flies were transferred into a fresh vial. The proportion of flies reaching the top of the vial within 20 seconds after knocking down was recorded. The results represent data collected from six replicates.

### Nucleolar volume measurement

The volume of Fibrillarin-stained nucleoli was determined based on Z-stack confocal images using the ‘3D object counter’ ImageJ plug-in. Objects on the edge of images were excluded, and threshold and size filters were automatically set. Data were obtained from three independent ovaries.

### Statistics

All experiments were conducted not less than 3 times independently. Statistical significance (*p*-value) was tested by applying paired *t*-tests with a 95% confidence interval or one-way ANOVA. All error bars represent the standard error of the mean (SEM). No statistical methods were used to predetermine sample size. Experiments were neither intentionally randomized nor intentionally ordered. Investigators were not blinded to allocation during experiments and outcome assessment.

## Data availability

Sequencing data generated during the current study will be made available in the NCBI Gene Expression Omnibus repository.

## Acknowledgments

We thank A. Teleman for reagents and antibodies; H. Ashe, A. Brand, F. Jiggins, O. Shimmi, the Vienna Drosophila Resource Center, and the Bloomington Drosophila Stock Center for fly reagents. A. Martinez-Arias for discussions and comments on the manuscript. TJS is a Herchel Smith Postdoctoral Fellow and KZAG is funded through an MRC PhD studentship. FKT is a Wellcome Trust and Royal Society Sir Henry Dale Fellow (206257/Z/17/Z) and is supported by the Human Frontier Science Program (CDA-00032/2018). For the purpose of Open Access, the author has applied a CC BY public copyright license to any Author Accepted Manuscript (AAM) version arising from this submission.

## Author contributions

JG and FKT conceived the idea and designed the experiments. JG, TJS and KZAG performed the experiments. FKT analyzed RNA sequencing data. JG, TJS and FKT wrote the manuscript.

## Competing interests

Authors declare no competing interests.

## Supplementary Figures

**Supplementary figure 1.**
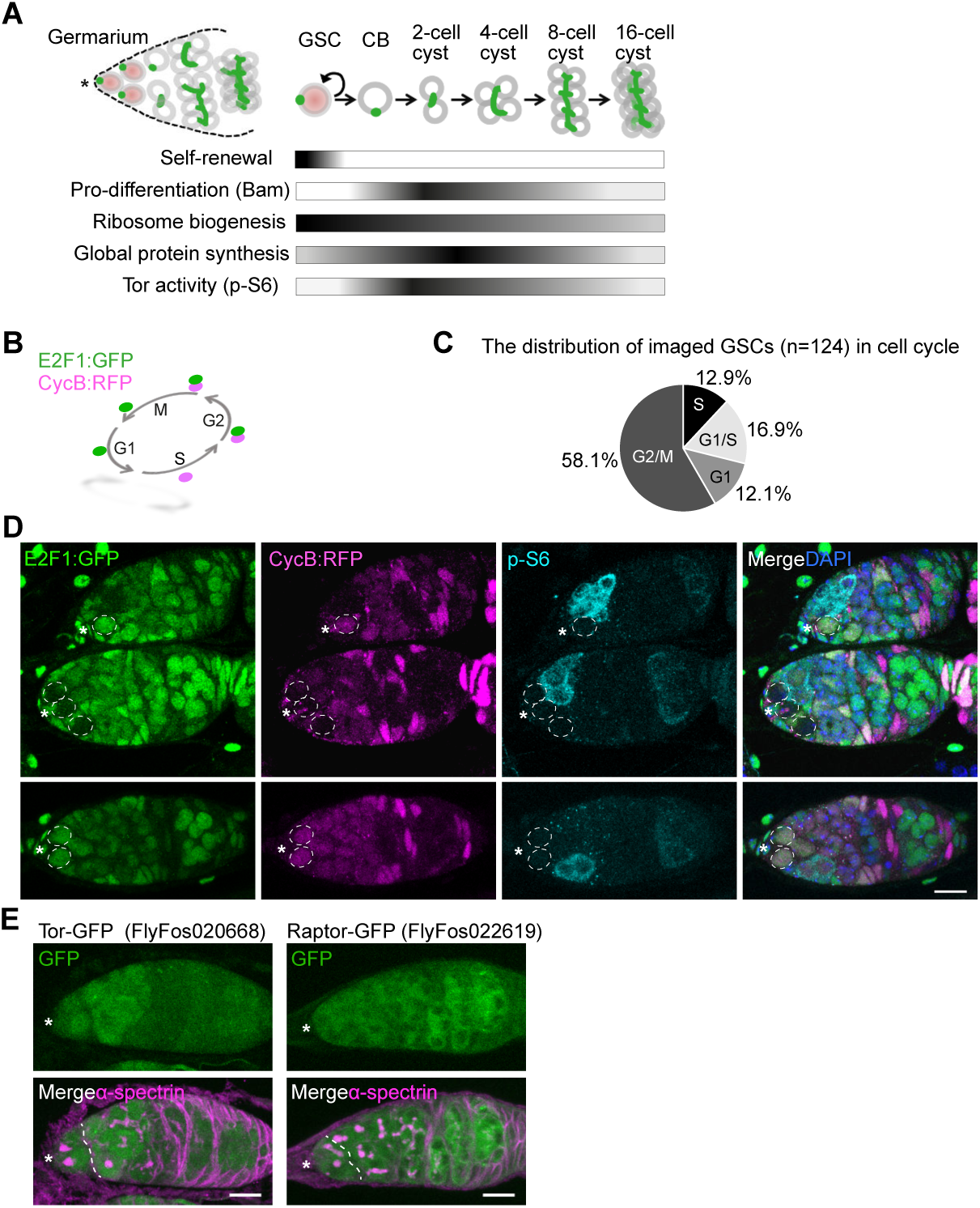
**(A)** Schematic of the structure of germarium and stages of GSC differentiation. Spectrosomes/fusomes in green. The asterisk indicates GSC niche. GSCs are marked by pink nuclei. **(B)** Schematic showing the rationale of the two-color fly FUCCI system. **(C)** Representative FUCCI germaria labeled with GFP (E2F1: GFP, green), RFP (CycB:RFP, magenta), p-S6 (Tor activity, cyan), and DAPI (nuclei, blue). Dashed circles mark GSCs. **(D)** Distribution of cell-cycle phase of imaged GSCs (n=124). **(E)** Representative image of germaria of transgenic Tor-GFP (left, FlyFos020668) and Raptor-GFP (right, FlyFos022619) flies labeled with α-spectrin (spectrosomes/fusomes, green) and GFP (fusion proteins, green). Dashed lines indicate the boundary between GSCs and differentiating cells. The asterisk indicates GSC niche (C, E). Scale bars, 10 μm (C, E).

**Supplementary figure 2.**
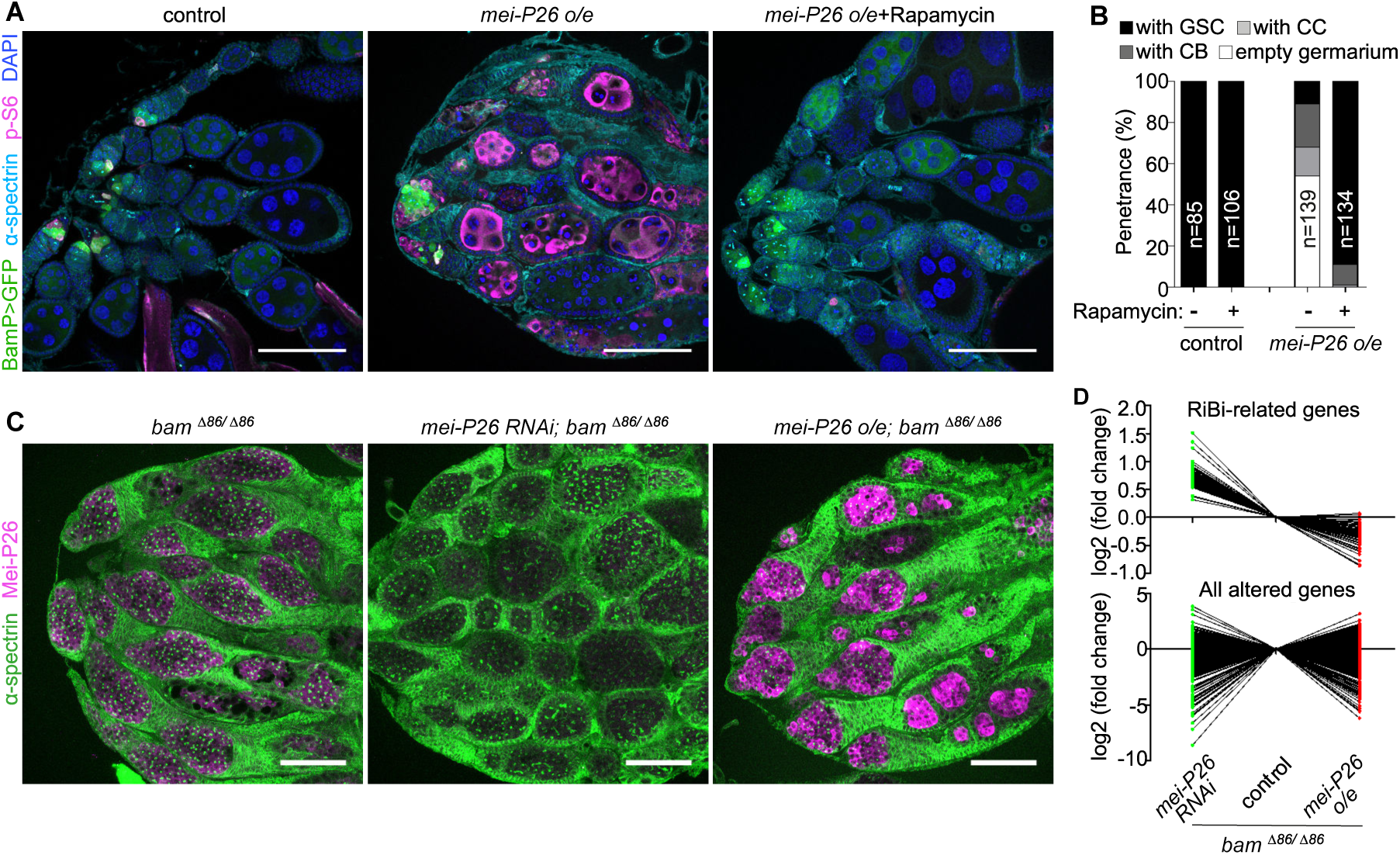
**(A)** Representative ovary images of control (*nos-gal4/+*) and Mei-P26 overexpression (o/e) (*nos-gal4/UASp-mei-P26)* flies with or without rapamycin feeding. Ovaries were stained with GFP (BamP>GFP, differentiating cells, green), p-S6 (Tor activity, magenta), α-spectrin (spectrosome/fusome, cyan), and DAPI (nuclei, blue). **(B)** Distribution of germarium phenotypes of control (*nos-gal4/+*) and Mei-P26 o/e (*nos-gal4/UASp-mei-P26)* with or without rapamycin feeding. **(C)** Representative ovary images of control (*nos-gal4/+*), *mei-P26* KD (*nos-gal4/UAS-mei-P26 RNAi)*, and *mei-P26* o/e (*nos-gal4/UASp-mei-P26)* in *bam*^*Δ86/ Δ86*^ background. Samples were stained with α-spectrin (green) and Mei-P26 (magenta). **(D)** RNA profiling of ovaries of control (*nos-gal4/+*), *mei-P26* KD (RNAi, *nos-gal4/UAS-mei-P26 RNAi)*, and *mei-P26* o/e (*nos-gal4/UASp-mei-P26)* in *bam*^*Δ86/ Δ86*^ background. Scale bars, 200 μm (A) or 100 μm (C).

**Supplementary figure 3.**
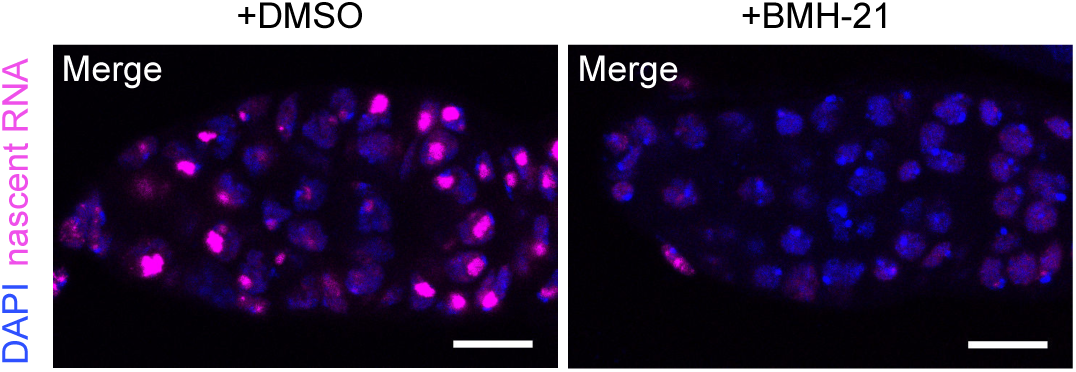
Representative confocal images of germaria treated with DMSO or BMH-21. Dissected ovaries were treated with DMSO or 100 μM BMH-21 (we estimate to be approximately equivalent to the concentration in the abdomen after injection in our experiments) for 3 hours, including 5-ethynyl uridine (EU) for the final 2 hours allowing visualization of nascent RNA. In the DMSO control most nascent RNA is generated in the nucleolus, and this is almost entirely abolished with BMH-21 treatment. Image is a max projection of 3 z planes. Scale bars, 10 μm.

**Supplementary figure 4.**
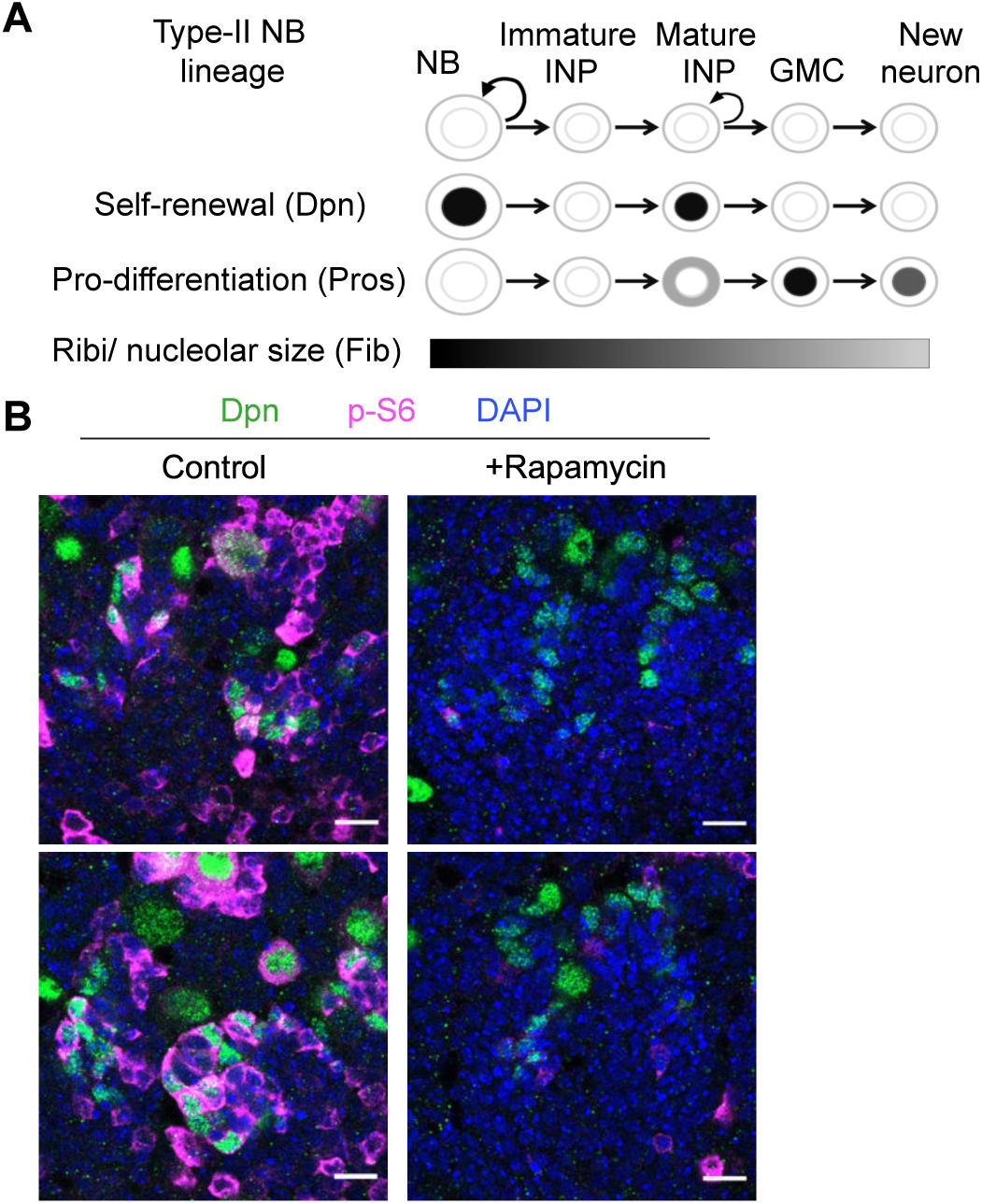
**(A)** Schematic showing different cell types and expression markers during type II NB differentiation. **(B)** Representative images of type II NB clusters after 20 minutes of *ex vivo* culture with or without 10 μM rapamycin. Samples are stained with Dpn (NBs and mature INPs, green), p-S6 (magenta) and DAPI (blue). Scale bars, 10 μm.

**Supplementary figure 5.**
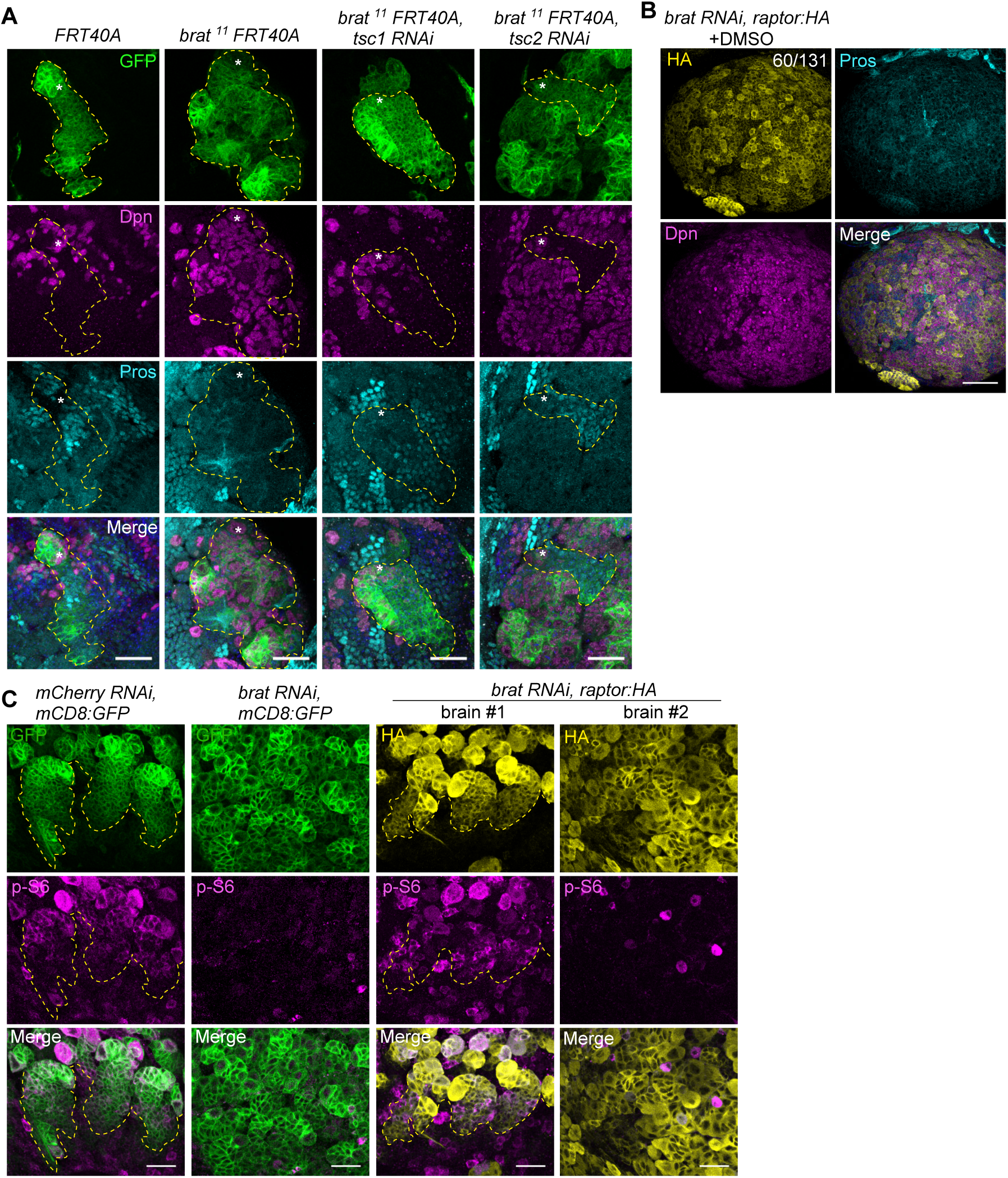
**(A)** Representative confocal images of MARCM clonal analysis of control (*FRT40A*), *brat* mutant (*brat*^*11*^ *FRT40A*), and *brat* mutant with *tsc1* or *tsc2* KD (*brat*^*11*^ *FRT40A, tsc1 RNAi or tsc2 RNAi*). Samples were stained with GFP (green), Dpn (magenta), Pros (cyan), and DAPI (blue). Dashed lines outline the clonal region. Asterisks indicate type II NBs within each clone. **(B)** Representative brain lobes of *brat* KD together with *raptor* overexpression (o/e) (*brat RNAi, raptor:HA*) flies (driven by *insc-GAL4*). Samples were stained with HA (yellow), Dpn (magenta), and Pros (cyan). The numbers in the upper panel indicate the penetrance of the phenotype. **(C)** Representative confocal images of type II NB lineages of control (*mCherry RNAi, mCD8:* GFP), *brat* KD (*brat RNAi, mCD8:* GFP), and *brat* KD with *raptor* o/e (*brat RNAi, raptor:HA*). Samples were stained with GFP/HA (green) and p-S6 (Tor activity, magenta). Dashed lines depict the border of *insc-GAL4* active regions (marked by GFP/HA). Scale bars, 50 μm (B) or 20 μm (C, A).

**Supplementary figure 6.**
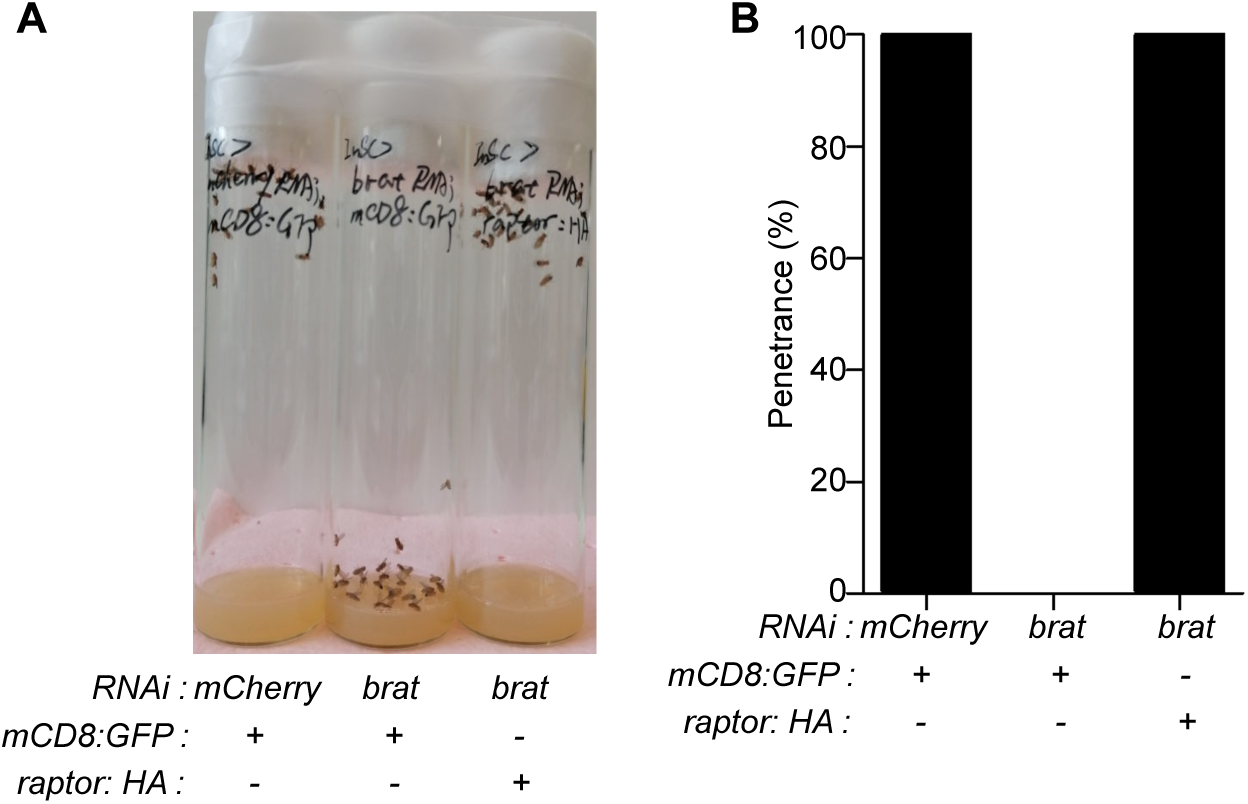
**(A)** Representative images of control (*mCherry RNAi, mCD8:GFP*), *brat* KD (*brat RNAi, mCD8:GFP*), and *brat* KD with *raptor* overexpression (o/e) (*brat RNAi, raptor:HA*) adult flies (driven by *insc-GAL4*). **(B)** Distribution of flies reaching the top of the vial in 20 seconds after knocking for controls (*mCherry RNAi, mCD8:GFP*), *brat* KD (*brat RNAi, mCD8:GFP*), and *brat* KD with *raptor* o/e (*brat RNAi, raptor:HA*).

**Supplementary figure 7.**
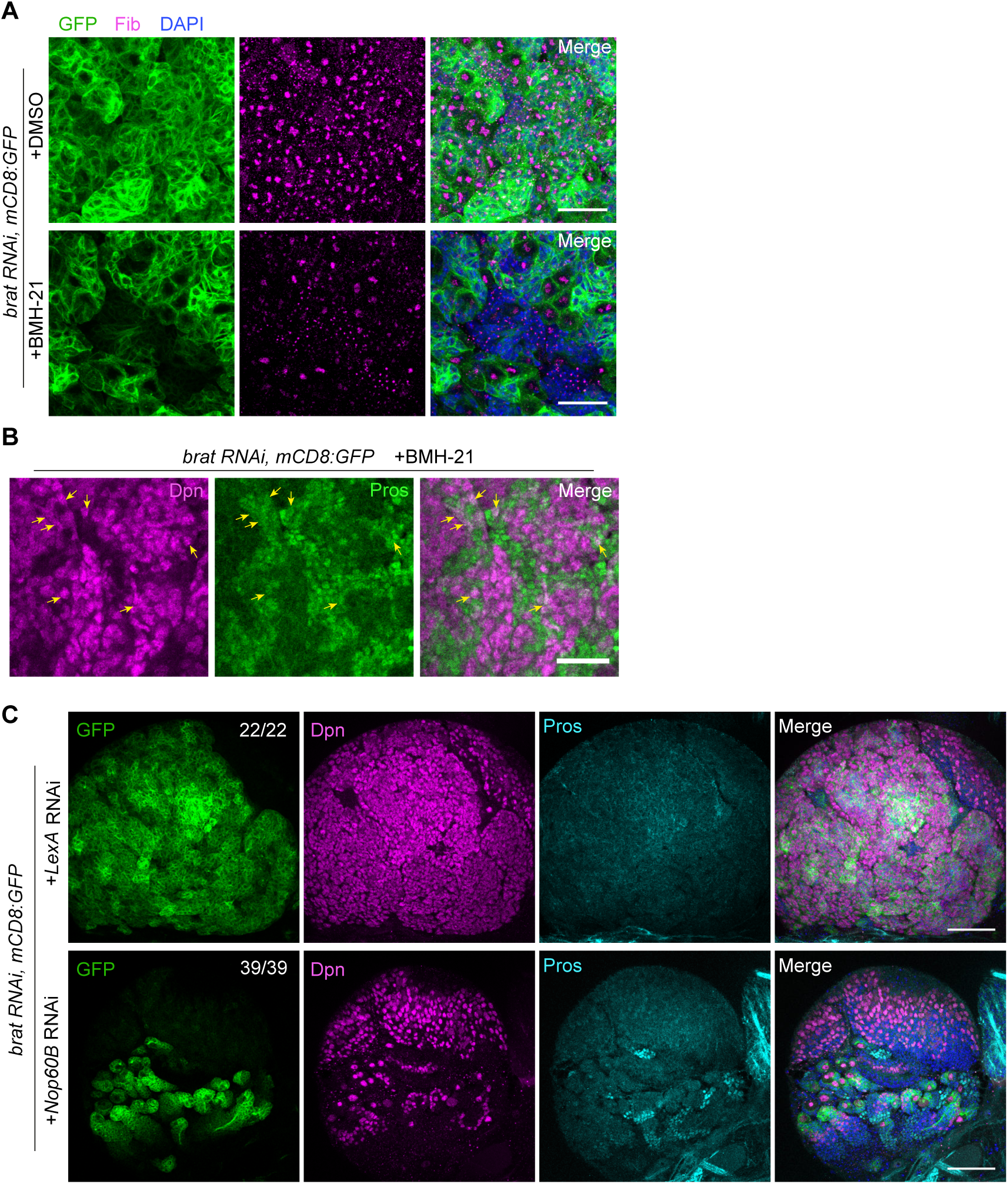
**(A)** Representative confocal images of *brat* KD (*brat RNAi, mCD8:GFP*) flies (driven by *insc-GAL4*) with or without BMH-21 feeding, stained with GFP (green), Fibrillarin (nucleoli, magenta), and DAPI (blue). **(B)** Representative confocal images of *brat* KD (*brat RNAi, mCD8:GFP*) flies (driven by *insc-GAL4*) with BMH-21 feeding, marked by Pros (magenta) and Dpn (green). Yellow arrows indicate the cells co-expressing Dpn and Pros markers. **(C)** Representative brain lobes of *brat* KD together with *LexA* RNAi or *nop60B* RNAi flies (driven by *insc-GAL4, UAS-mCD8:GFP*). Samples were stained with Dpn (magenta), Pros (cyan), and DAPI (blue). The numbers indicate the penetrance of the phenotype. Scale bars, 50 μm (C), 20 μm (A) or 10 μm B).

**Supplementary table 1**. Results of RNA-seq analysis in the *bam*^*Δ86*^ mutant background.

**Supplementary data-video 1**. Representative movie of adult climbing assay. From left to right: *mCherry* RNAi control, *brat* RNAi and rescue *brat* RNAi with *raptor:HA*.

## Notes

### Competing Interest Statement

The authors have declared no competing interest.

